# Global Stability and Tipping Point Prediction of the cell fate decision making model

**DOI:** 10.1101/2024.12.17.629061

**Authors:** Li Xu, Jin Wang

**Affiliations:** State Key Laboratory of Electroanalytical Chemistry, Changchun Institute of Applied Chemistry, Chinese Academy of Sciences, Changchun, Jilin, 130022, P.R.China; Department of Chemistry and of Physics and Astronomy, State University of New York at Stony Brook, Stony Brook, NY 11794-3400, USA

## Abstract

This study investigates global stability and tipping point prediction in cell fate decision-making systems through non-equilibrium landscape flux theory. We demonstrate that cell fate dynamics are governed by the interplay between potential landscape and curl flux, where the landscape guides systems toward stable states while curl flux mediates transitions between them.

Our analysis reveals that non-zero curl flux generates irreversible dominant pathways between multi-stable states. We identify several quantitative measures for transition prediction, including barrier heights, kinetic switching times, entropy production rates, and average flux. We introduce novel non-equilibrium early warning indicators based on time irreversibility of cross-correlations (Δ*C*), average flux, and entropy production rate. These indicators exhibit significant changes near bifurcations, enabling transition prediction before state stability loss, with superior predictive capability compared to traditional critical slowing down theory.

The rotational nature of curl flux is shown to destabilize attractor states, providing a dynamical foundation for phase transitions in cell differentiation, reprogramming, and transdifferentiation processes. These findings advance our understanding of non-equilibrium dynamics in cell fate decisions and offer practical implications for stem cell research and regenerative medicine, potentially enabling more precise therapeutic strategies in stem cell applications.

## I. INTRODUCTION

Cell fate decision making, wherein cells determine their specific developmental or functional identity, remains a fundamental challenge in biological systems. The processes of cell reprogramming and transdifferentiation present significant opportunities for stem cell applications^1^, with transdifferentiation occurring through two distinct pathways^2^. The first pathway involves the reprogramming of mature cells into a progenitor-like state through the action of specific transcription factors and signaling molecules^3^. These reprogrammed progenitor cells can subsequently be directed toward desired cell types for therapeutic applications^4^. This approach shows particular promise in regenerative medicine for tissue repair and replacement^5^.

The second pathway, as illustrated in Figure 1, involves direct transdifferentiation that bypasses the progenitor stage, offering a more efficient approach for generating specific cell types. This direct conversion method has broad therapeutic applications, including the transformation of fibroblasts into functional neurons for neurodegenerative diseases, conversion of liver cells into pancreatic beta cells for diabetes treatment, and modification of cardiac cells for heart repair^6–10^.

**FIG. 1.**
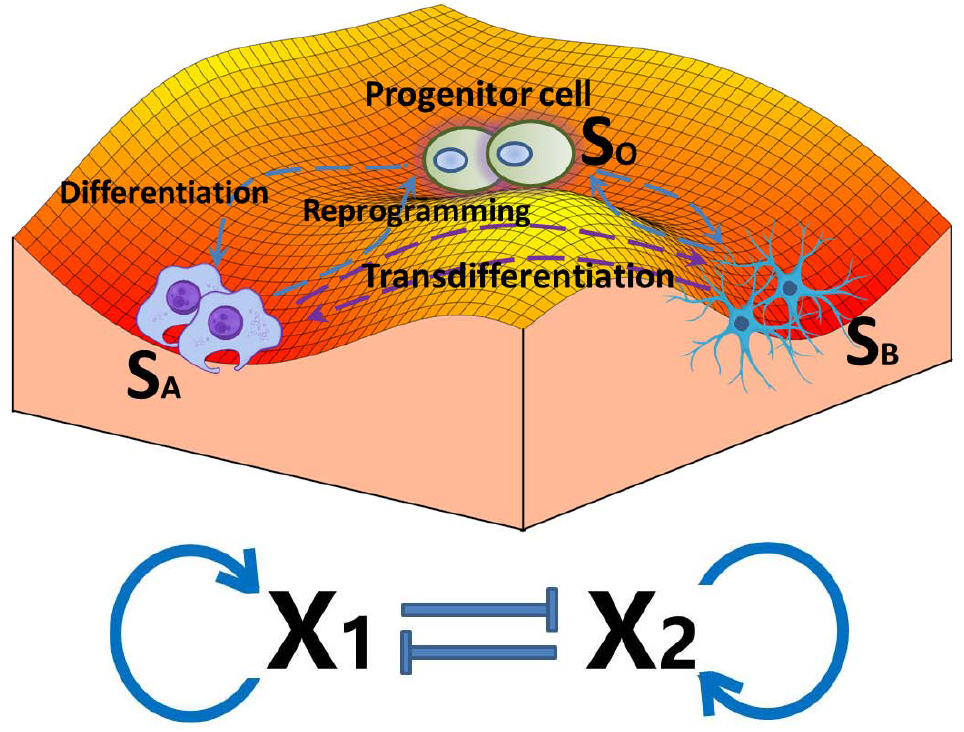
Scheme of transdifferentiation, reprogramming and differentiation: *X*_1_ and *X*_2_ are mutually inhibitory and self-activating gene regulatory networks.

A significant challenge in controlling transdifferentiation lies in the prediction and direction of cell fate decisions^11^. Recent advances have revealed important “early warning signals” through transcriptomic and epigenetic modifications^12^. Understanding these molecular signals is essential for decoding the mechanisms that govern cell fate decisions during transdifferentiation, ultimately advancing the development of targeted cell-based therapies^13^.

Biological systems frequently undergo abrupt regime shifts, transitioning between distinct stable states with potentially significant implications^12^. Critical slowing down (CSD) provides a theoretical framework for predicting these tipping points in complex systems^14^. As a system approaches a critical transition, CSD manifests through increasingly sluggish responses to perturbations, characterized by two fundamental indicators: enhanced autocorrelation, reflecting increased memory of previous states, and extended relaxation time required to return to equilibrium^15^.

While CSD has proven effective in predicting tipping points, its application has largely been confined to one-dimensional data analysis, presenting limitations in understanding complex biological systems^16^. This constraint is particularly relevant in cell fate decision making processes, where multiple molecular and cellular components interact in nonlinear ways. To address this limitation, our study explores the application of differences in cross-correlation analysis to multi-dimensional, nonlinear cell fate decision making during transdifferentiation and reprogramming processes. This approach aims to enhance the prediction accuracy of regime shifts in cellular systems and deepen our understanding of the underlying mechanisms governing cell fate transitions.

Current modeling approaches frequently focus on isolated differentiation stages or lack comprehensive analysis capabilities for early warning signals in cell fate decision making processes^17^. To address these limitations, we implement a generalized fate decision-making model incorporating ordinary differential equations that capture essential components influencing cell state transitions and gene expression dynamics^18^.

We employ potential landscape and flux theory to investigate early warning signals for cell transdifferentiation^7^. Our analysis identifies three key early warning indicators: entropy production rate, average flux, and the average difference in cross-correlations between forward and backward time directions. Notably, both the average curl flux and entropy production rate reach maximum values prior to bifurcation or dynamical phase transition points when regulatory mechanisms or interaction strengths vary. This suggests that rotational curl flux serves as a dynamic origin, while entropy production rate functions as a thermodynamic origin for bifurcation/catastrophe transitions in non-equilibrium cell development systems^19^.

Our approach enables identification of tipping points - critical parameter values where the cell fate landscape undergoes fundamental reorganization. Significantly, the time-reversal symmetry-breaking characteristics, which reflect dynamical and thermodynamical warning signals, can be detected earlier than traditional indicators based on nonlinearity, dynamics, and bifurcation theory^12^. The irreversibility observed in time series data strongly correlates with these dynamical and thermodynamic quantities, providing a practical method for transition forecasting. This predictive capability has substantial implications for stem cell engineering, regenerative medicine, and disease modeling applications^13^.

## II. METHODS

### A. Cell fate decision making Model

We investigated a pivotal gene regulatory circuit module governing cell differentiation and development, which encompasses two genes crucial for cell fate determination. Illustrated in Figure 1 as *X*_1_ and *X*_2_, these genes engage in mutual inhibition and self-activation within this regulatory circuit^6–9^. The expression levels of genes *X*_1_ and *X*_2_ are denoted as *X*_1_ and *X*_2_ respectively, capturing their time-dependent dynamics. To elucidate the expression dynamics of these genes, we describe the cell fate decision-making circuit through a set of two ordinary differential equations:

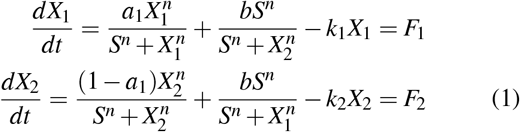

The parameters interpretations and default values of the parameters are given in Table I^6–9^. The sum of *a*_1_ and *a*_2_ is

**TABLE I.** Parameters interpretation and default values^6–9^.

| Symbol | Ecological interpretation | Default value |
| --- | --- | --- |
| $a_1$ | the self-activation strength of gene $X_1$ | — |
| $a_2$ | the self-activation strength of gene $X_2$ | — |
| $b$ | the mutual inhibition strength | — |
| $S$ | the Hill function incorporating threshold parameter | 0.5 |
| $k_1$ | the first-order degradation rate governing gene $X_1$ | 1.0 |
| $k_2$ | the first-order degradation rate governing gene $X_2$ | 1.0 |
| $n$ | the Hill function cooperative parameter | 4.0 |

1.The regulation of *a*_1_ and *a*_2_ can induce the process of cell transdifferentiation.

### B. Landscape and flux theory for non-equilibrium cell fate decision making systems

The landscape and flux theoretical framework has been established and applied across various fields^6–9,20–24^. This framework offers a robust methodology for investigating the global stability and bifurcations inherent in non-equilibrium dynamical systems.

Biological systems commonly exhibit fluctuations^7,21,25^. The dynamics upon fluctuating environments can be expressed as: 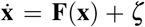 . Here, **F**(**x**) represents the deterministic force, with **x** denoting the vector encompassing various variables such as concentrations in state space. The term *ζ* signifies a Gaussian fluctuating force, characterized by an autocorrelation function given as *< ζ* (**x**, *t*)*ζ* (**x**, 0) *>*= 2**D**(**x**)*δ* (*t*), where **D**(**x**) denotes the diffusion coefficient matrix. We define **D**(**x**) = *D***G**(**x**), where *D* represents the diffusion coefficient, reflecting the magnitude of noise strength, and **G** signifies the scaled diffusion matrix, describing anisotropy and inhomogeneity. For simplicity, in this study, we consider **G**(**x**) as a unit matrix.

Instead of traversing unpredictable stochastic evolutionary trajectories governed by the Langevin equation, we can explore the corresponding Fokker-Planck diffusion equation^26–28^, governing the evolution of probability *P*(**x**, *t*), which exhibits linearity, determinism, and predictability: *∂P/∂t* = −∇ · (**F***P* − *D***G**(**x**)∇*P*). The probability flux **J** is mathematically defined as **J** = **F***P*−*D*∇*P*. It is demonstrated that the driving force for the system dynamics can be expressed as **F** = −*D*∇*U* + **J***ss/Pss*^6,7,20–23^, where *U* denotes the nonequilibrium potential landscape, defined as *U* = −ln*P*_*ss*_, associated with the steady-state probability distribution *P*_*ss*_, and **J**_*ss*_ denotes the steady-state probability flux. At steady state, ∇ · **J** = 0, implying that the steady-state probability flux can either be zero, constant, or exhibit rotational (curl-like) behavior. Equilibrium state, characterized by zero flux, signifies detailed balance where there is no net input or output. As the nonzero flux represents net flow into or out of the system, the flux magnitude serves as a metric quantifying the extent of detailed balance deviation from equilibrium.

The non-equilibrium open systems exchange energy, materials or information with the environments. The system entropy evolution in time can be decomposed to the entropy production rate and heat dissipation rate^20,22,29–33^: 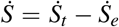, where 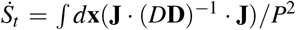 represents the total entropy production rate of the system and the envioronment, which is non-negative. Meanwhile, 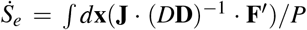 signifies the heat dissipation rate, which can be either positive or negative. This rate quantifies the flow of entropy from the environment to the non-equilibrium system. Notably, in steady state conditions, the entropy production rate coincides with the heat dissipation rate^20,22,29,30^. Consequently, the entropy production rate serves as a comprehensive thermodynamic descriptor for the non-equilibrium system.

To understand better the underlying dynamical process, one can find the most likely paths between stable states by the path integral method. This method calculates the probability of a path from an initial state, denoted by **x**_*i*_ at time *t* = 0, to a final state, denoted by **x** _*f*_ at a later time *t*. The probability is determined by the Lagrangian, *L*(**x**(*t*)), of the system and the action, *A*(**x**), associated with the path. The path integral formula is shown as^18,23^:

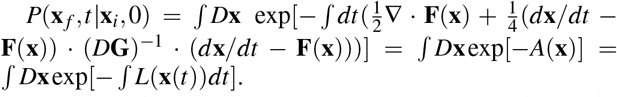

Here, *D***x** denotes the summation over all feasible paths connecting the initial state, **x***i* at time zero, to the final state **x** *f* at time *t*^18,23^. By minimizing the weight function, or the action represented by *A*(**x**), we can find the most probable trajectories as the dominant paths among these potential paths.

## III. RESULTS

Stem cell research hinges on our ability to control how cells swtich between different states. Non-equilibrium landscape flux theory offers a powerful framework to explore the dynamics of cell fate decision making model under both finite and zero fluctuations. In this system, genes *X*_1_ and *X*_2_ form a unique regulatory network, where they both inhibit each other and activate themselves. By manipulating the self-activation strength of these genes, we can potentially induce transdifferentiation, the process by which a cell converts directly into another cell type. There are two main pathways for transdifferentiation shown in Figure 1: 1.Transdifferentiation with reprogramming, A differentiated cell (*S*_*A*_) is first reprogrammed to a less specialized, intermediate pluripotent state (*S*_*O*_) with greater developmental potential. Then, it can be subsequently induced and re-differentiated into another lineage differentiated cell type *S*_*B*_ (the blue arrow lines); 2.Direct transdifferentiation, a differentiated cell (*S*_*A*_) is directly converted into another cell type (*S*_*B*_) without going through an intermediate stage (the purple arrow lines). Cell type transitions can be achieved via various mechanisms including induction, gene regulation, and stochastic fluctuations^6–9,34^.

### A. Transdifferentiation with Reprogramming

We investigated the bifurcation phenomena inherent in the gene circuit responsible for cell differentiation across various regulatory parameter settings (the mutual inhibition strength *b* = 0.21) shown in Figure 2A. The process of cell transdifferentiation is characterized by four saddle-node bifurcations. There are three stable states in the phase diagram. *S*_*A*_ represents the lineage-specific stable state with lower expression of gene *X*_1_ and higher expression of gene *X*_2_. *S*_*B*_ represents the lineage-specific stable state with lower expression of gene *X*_2_ and higher expression of gene *X*_1_. *S*_*O*_ represents the pluripotent stable state with intermediate expression of gene *X*_1_ and *X*_2_. Through the solving of the Fokker-Planck diffusion equation, the probability distribution of the system at steady state can be obtained, enabling the derivation of the potential landscape of the system via the *U* = −ln *P*_*SS*_. We show the three-dimensional potential landscapes versus *a*_1_ with the diffusion coefficient *D* = 0.005 and *b* = 0.21 in Figure 3A. When the self-activation strength *a*_1_ is relatively small and the self-activation strength parameter *a*_2_ is relatively high, the system stabilizes in a differentiated cell state *S*_*A*_ (*a*_1_ = 0.1). With an increase in the self-activation strength parameter *a*_1_ and a concurrent decrease in *a*_2_, following the initial saddle-node bifurcation, a new stable stem cell state *S*_*O*_ emerges alongside the differentiated cell state *S*_*A*_ (*a*_1_ = 0.3, 0.38). Subsequently, as *a*_1_ continues to rise and *a*_2_ decreases further, upon the second saddle-node bifurcation, the *S*_*A*_ state vanishes, leaving only the pluripotent state *S*_*O*_ (*a*_1_ = 0.50). Further increment of *a*_1_ and reduction of *a*_2_ lead to the occurrence of the third saddle-node bifurcation, resulting in the emergence of another cell differentiation state *S*_*B*_, coexisting with *S*_*O*_ (*a*_1_ = 0.70). Ultimately, with a significant increase in *a*_1_ and a substantial decrease in *a*_2_, the system experiences the fourth saddle-node bifurcation, causing the disappearance of the *S*_*O*_ state and leaving behind only the differentiated state *S*_*B*_ (*a*_1_ = 0.90) in Figure 2A. The parts with yellow-green background are the bistable region.

**FIG. 2.**
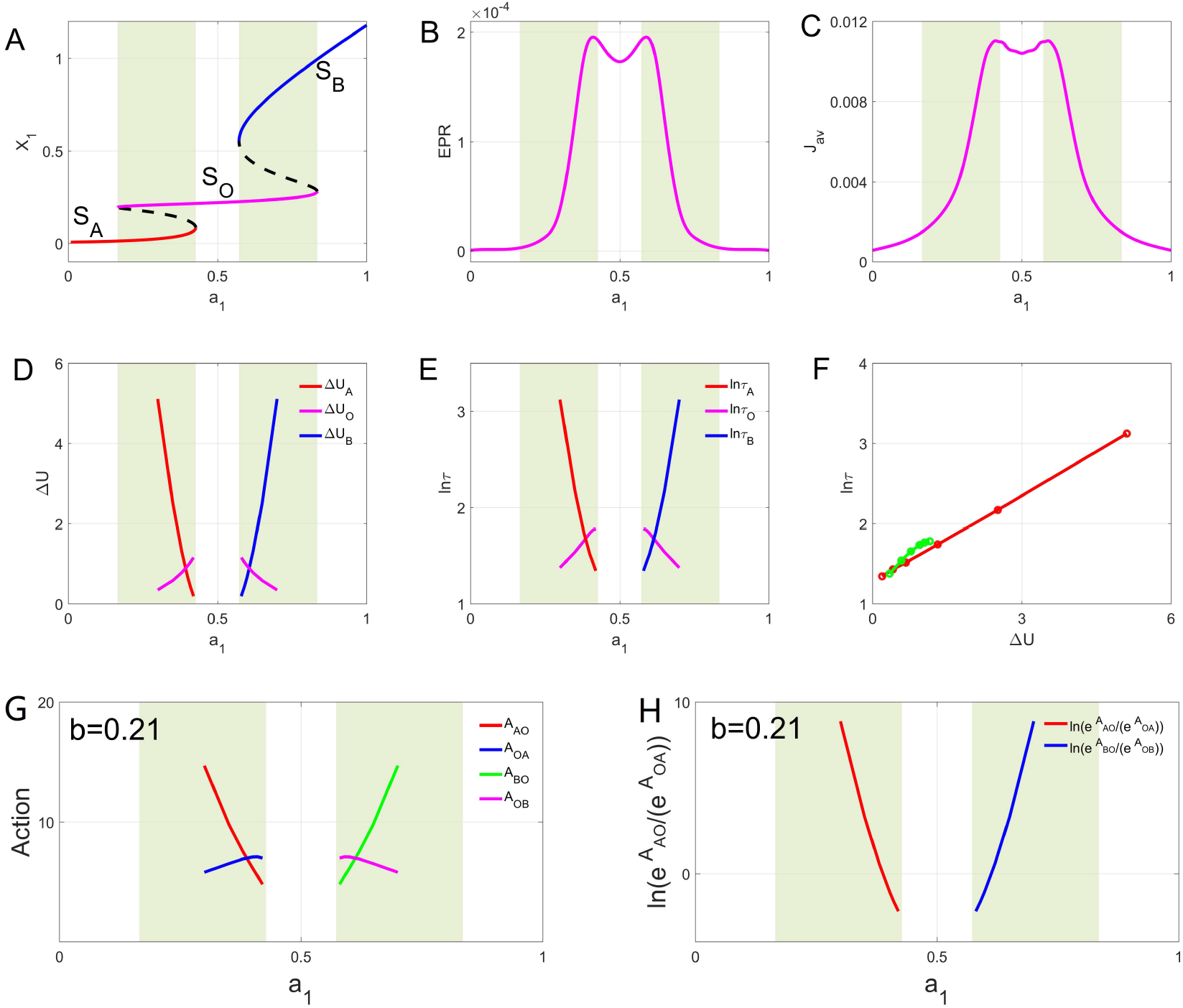
A: The phase diagram of two mutual inhibition and self-activation genes *X*_1_ and *X*_2_ with increasing the self-activation strength *a*_1_ giving arise to saddle node bifurcations (*b* = 0.21). B: The entropy production rate (*EPR*) versus *a*_1_. C: The average curl flux *J*_*av*_ versus *a*_1_. D: The barrier heights Δ*U* versus *a*_1_. E: The logarithm of the mean first passage time versus *a*_1_. F: The logarithm of the mean first passage time versus the barrier heights Δ*U* . G: The action *A*_*AO*_ of the probability of the dominant path from *S*_*A*_ to *S*_*O*_ and the action *A*_*OA*_ of the probability of the dominant path from *S*_*O*_ to *S*_*A*_ versus *a*_1_. The action *A*_*BO*_ of the probability of the dominant path from *S*_*B*_ to *S*_*O*_ and the action *A*_*OB*_ of the probability of the dominant path from *S*_*O*_ to *S*_*B*_ versus *a*_1_. H: The logarithm of the action *A*_*AO*_ divided by the action *A*_*OA*_ versus *a*_1_. The logarithm of the action *A*_*BO*_ divided by the action *A*_*OB*_ versus *a*_1_. The parts with yellow-green background are the bistable region.

**FIG. 3.**
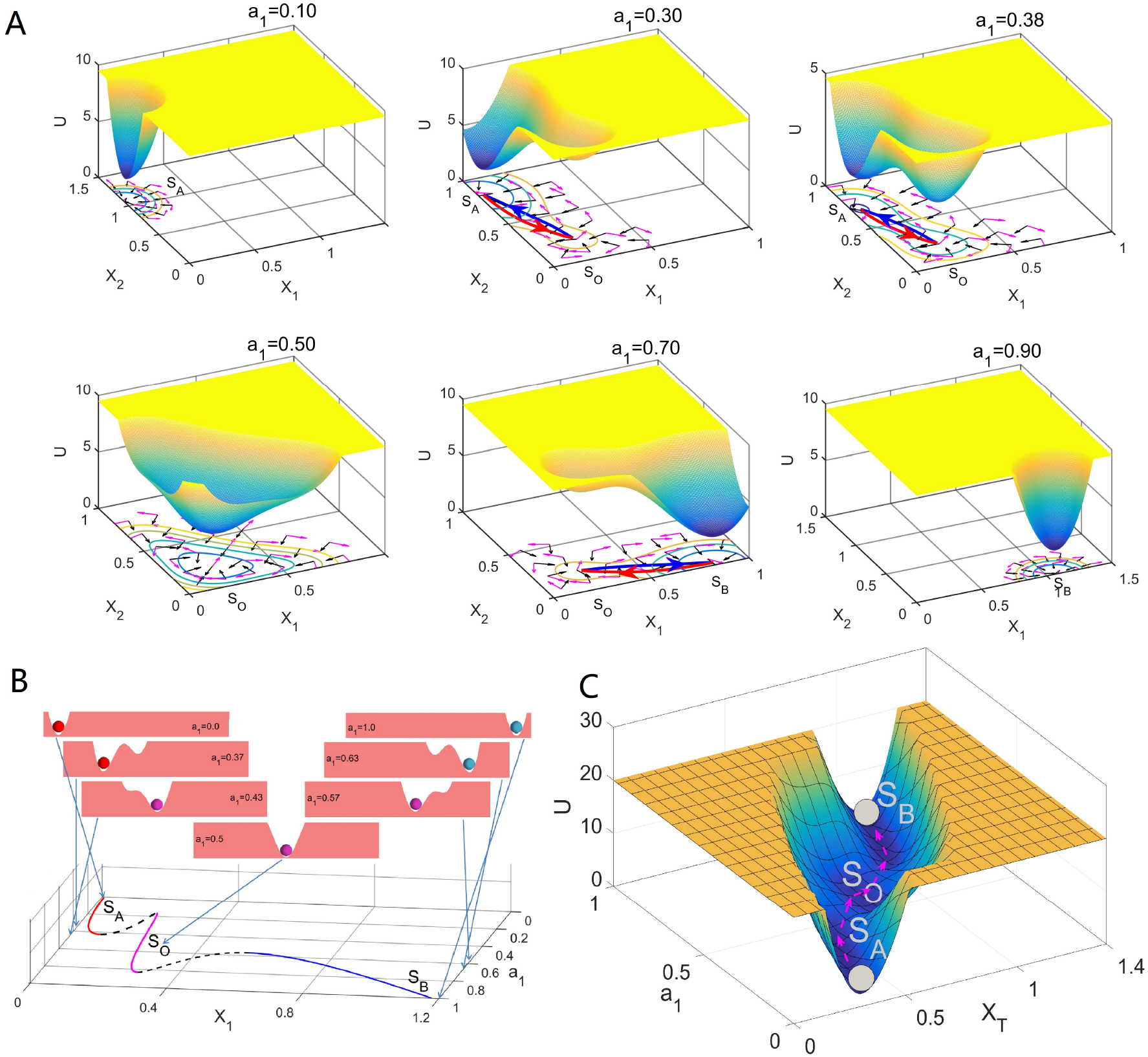
A: The potential landscapes versus *a*_1_ with the diffusion coefficient *D* = 0.005 (*b* = 0.21). The pathways on potential landscapes. The red lines represent the dominant population paths from the *S*_*A*_(*S*_*B*_) state to the *S*_*O*_ state. The blue lines represent the dominant population paths from the *S*_*O*_ state to *S*_*A*_(*S*_*B*_) state. B: The quantitative one-dimensional population potential landscapes by the projection versus *a*_1_ are on the top. The phase diagram is on the bottom. C: The continuous potential landscape *U* versus the *a*_1_ and *X*_*T*_ with integral. The projection line is *X*_2_ = −*X*_1_ + 2 with the coordinate transformation 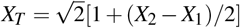.

We depict the potential landscape *U* projected onto the line *X*_2_ = −*X*_1_ + 2 and visualize it in Figure 3B with the mutual inhibition strength *b* = 0.21. This potential landscape serves as a quantitative assessment of the probability distribution within cell fate decision making model. The one-dimensional potential landscape, while quantitatively accurate, bears qualitative resemblance to the Waddington model or the metaphorical representation of a ball resting in a valley to depict ecosystem stability^35,36^. This visualization aids in understanding steady-state displacements using the conceptual framework of a “ball in the valley.” Under small fluctuations, the system deviates momentarily from its steady state. When positioned on a hillside, the system tends to return to stability due to the topographical features resembling a valley. Consequently, minor fluctuations allow the ball to return to its stable position at the valley bottom. However, if fluctuations intensify, the ball may traverse the unstable saddle point’s ridge and swith into an adjacent stable valley, leading the system into another steady-state region^12,15,37,38^. The ball’s location at the valley bottom signifies system stability, characterizing the strength of the attractor within the system. Figure 3C illustrates a three-dimensional continuous potential landscape *U* versus *a*_1_ projected on<u>t</u>o the line *X*_2_ = −*X*_1_ + 2, with the transformation 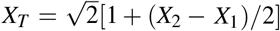 . This representation also bears qualitative resemblance to the marble-in-a-cup model, a metaphor often used to conceptualize the stability^35,36^. This depiction encapsulates a potential landscape quantified by probability distributions.

Figure 2BC shows the entropy production rate (*EPR*) and the average curl flux *J*_*av*_ versus the self-activation strength *a*_1_ with the mutual inhibition strength *b* = 0.21. We can see that *EPR* and *J*_*av*_ have the same trends and both have *M* shape. The two lines increase approaching its peak value first and then decreases and repeat this process once. The observed behavior is attributed to the distribution of curl flux. For lower values of the self-activation strength *a*_1_, only one stable differentiated cell state *S*_*A*_ exists, and the curl flux is distributed around a single stable attractor (Figure 3). However, as the self-activation strength *a*_1_ approaches its dynamical phase transition or bifurcation tipping point, a new stable stem cell state *S*_*O*_ begins to emerge. This leads to a broader and more intense distribution of curl flux within the state space (Figure 3). In addition to the flux revolving around the emerging stem cell state *S*_*O*_, there is also flux between *S*_*O*_ and *S*_*A*_ (Figure 3). The additional curl flux between the steady states contributes to an increase in the average flux *J*_*av*_ and *EPR*, and concurrently, the entropy production rate (*EPR*), closely linked to flux, also reaches a maximum. As the self-activation strength *a*_1_ continues to increase, the stable attractor *S*_*A*_ exhibit the decrease of its stability. With the disappearance of the *S*_*A*_ state, the two attractors change to a single stable attractor system, resulting in a reduced distribution of curl flux within the state space, although still greater than when only *S*_*A*_ exists alone (Figure 3). Consequently, both *J*_*av*_ and *EPR* experience a slight decrease, yet their values remain considerably higher than those observed when the *S*_*A*_ state is present. With further increases in the self-activation strength *a*_1_, a symmetrical change process unfolds within the system. In non-equilibrium systems, the interplay between the potenital landscape and the curl flux dictates the system’s dynamics. The landscape, characterized by the potential gradient, intrinsically stabilizes specific states. Conversely, the curl flux, with its rotational nature, exerts a destabilizing force, propelling the system away from these stable states (point attractors). This interplay is evident in our observations: increasing the curl flux leads to the destabilization of the original attractor and the emergence of two new attractors. In simpler terms, the flux alters the landscape topography, influencing the formation of phases and triggering bifurcations. Consequently, the curl flux, along with its associated entropy production rate (EPR), serves as a quantitative measure for driving phase transitions or bifurcations in non-equilibrium systems. We further observe that the fluxes rotate around individual stable states and spread between them. Higher flux magnitudes indicate greater dissipation, as measured by the EPR, which facilitates the communication or “association” between different stable attractors.

The pathways are on bottom of potential landscapes shown in Figure 3A in the two-stable-state-coexistence regions (yellow-green background regions in Figure 2A). In Figure 3A, the red lines represent the dominant population paths from the *S*_*A*_(*S*_*B*_) state to the *S*_*O*_ state. The black lines represent the dominant population paths from the *S*_*O*_ state to *S*_*A*_(*S*_*B*_) state. The purple arrows denote the steady-state probability fluxes that direct the dominant pathways, deviating from the anticipated steepest descent path, typically expected to traverse through the saddle point solely based on the potential landscape. Conversely, the black arrows depict the negative gradients of the potential landscapes. We observe distinct dominant pathways for cell reprogramming from the differentiated stable cell state *S*_*A*_(*S*_*B*_) to the pluripotent stable state *S*_*O*_ and for cell differentiation from the *S*_*O*_ state back to the *S*_*A*_(*S*_*B*_) state. This divergence in pathways contrasts with the expected scenario in equilibrium conditions where pathways remain identical. Notably, the discrepancy between the red and black pathways shows the inherent differences between cell differentiation and reprogramming pathways, rendering the transformation “irreversible”. The irreversibility observed in the state transitions is attributed to the existence of non-equilibrium curl flux, which serves as a defining characteristic of the non-equilibrium system.

Figure 2D depicts the barrier heights of the potential landscape as a function of *a*_1_. The light green background delineates the bistable region of the system. We define Δ*U*_*A*_ = *U*_*S*_ − *U*_*A*_, Δ*U*_*O*_ = *U*_*S*_ − *U*_*O*_, and Δ*U*_*B*_ = *U*_*S*_ − *U*_*B*_, where *U*_*S*_ represents the potential value of the saddle point, *U*_*A*_ signifies the potential value of the differentiated cell state *S*_*A*_, *U*_*B*_ corresponds to the differentiated cell state *S*_*B*_, and *U*_*O*_ denotes the potential value of the stem cell state *S*_*O*_. In Figure 2D, it is evident that as *a*_1_ increases, the barrier height Δ*U*_*A*_ of the differentiated cell state *S*_*A*_ decrease, while the barrier height Δ*U*_*B*_ of the differentiated cell state *S*_*B*_ increases. Additionally, the barrier height Δ*U*_*O*_ of the stem cell state *S*_*O*_ initially increases and then decreases with the rise in *a*_1_. The barrier height serves as an indicator of the stability of the steady state. The intersection of the barrier heights signifies an equal weighting of their respective stabilities, indicating that the two attractors are equally preferred.

Figure 2E depicts the logarithm of the mean first passage time (MFPT) versus *a*_1_. Here, *τ*_*A*_(*τ*_*B*_) represent the MFPTs to escape from the differentiated cell states *S*_*A*_(*S*_*B*_), respectively, while *τ*_*O*_ denotes the MFPT to escape from the stem cell state *S*_*O*_. The kinetic speed or kinetic time can be quantified by the MFPT from one state to another. The MFPT can be derived from the following equation^26^: **F** · ∇*τ* + *D*∇*τ* · **D** · ∇*τ* = −1. Our analysis reveals that increasing *a*_1_ leads to a decrease in ln *τ*_*A*_, an increase in ln *τ*_*B*_, and a non-monotonic trend in ln *τ*_*O*_, with an initial increase followed by a decrease. Notably, the intersection of the ln *τ*_*A*_ (ln *τ*_*B*_) and ln *τ*_*O*_ curves corresponds to the point where the weights of states *S*_*A*_(*S*_*B*_) and *S*_*O*_ become equal. This is the first order phase transition also coincides with the intersection of the barrier heights and the maximum point of *J*_*av*_ and *EPR*, which occurs at approximately *a*_1_ = 0.4. Figure 2F explores the relationship between the mean first passage time and the barrier heights on the potential landscape. Interestingly, the plot reveals a near-exponential correlation between these two quantities, suggesting that *τ* ∼ *e*^Δ*U*^ . This implies that the escape time from a cell state increases exponentially with the height of the landscape barrier separating it from the next state. In simpler terms, cells require a longer time to overcome a more substantial landscape barrier during transitions between states.

Dominant paths can also reveal shifting stability of potential landscapes. Identifying the most probable transition paths between stable states is crucial for understanding cell fate decisions in our model. The dominant path, characterized by the largest transition probability, offers valuable insights into these pathways. Figure 2G quantifies the dominant path probability using the action, *A*(**x**). Here, *A*_*AO*_(*A*_*BO*_) represents the action associated with the dominant path from state *S*_*A*_(*S*_*B*_) to state *S*_*O*_, while *A*_*AO*_ and *A*_*OB*_ denotes the action for the dominant path from *S*_*O*_ to *S*_*A*_(*S*_*B*_). Importantly, a higher action value signifies a lower dominant path probability, as this dominant path probability is proportional to exp[ −*A*(**x**)]. As expected, our results in Figure 2G demonstrate that *A*_*AO*_ and *A*_*OB*_ decrease, while *A*_*BO*_ and *A*_*OA*_ increase with increasing self-activation strength (*a*_1_). Figure 2H further corroborates this by showing an increase in the logarithm of the dominant path probability from *S*_*A*_ to *S*_*O*_ relative to the dominant population path from *S*_*O*_ to *S*_*A*_ as *a*_1_ increases. These observations collectively suggest that state *S*_*A*_ becomes less stable, while state *S*_*O*_ becomes more stable with increasing *a*_1_. Consequently, transitions from *S*_*O*_ to *S*_*A*_ become progressively more challenging, while transitions from *S*_*A*_ to *S*_*O*_ become more facile. The analysis of dominant path probabilities reveals a mirroring trend for transitions from state *S*_*B*_ to state *S*_*O*_. The logarithm of the dominant path probability from *S*_*B*_ to *S*_*O*_ relative to the dominant population path from *S*_*O*_ to *S*_*B*_ decreases with increasing self-activation strength (*a*_1_).

We explore an approach to quantify the non-equilibrium nature of the cell fate decision making system by analyzing the temporal irreversibility of its time series data. Here, we leverage the stochastic Langevin equation to simulate the long-term trajectories of variables *X* and *Y* as the noise-induced attractor fluctuates between states *S*_*A*_, *S*_*O*_ and *S*_*B*_. To quantify the time irreversibility, we employ the concept of cross-correlation functions. The forward in time cross-correlation function, denoted by *C*_*XY*_ (*τ*) = (*X* (0)*Y* (*τ*)) , is defined as the ensemble average of the product of *X* and *Y* , where *τ* represents the time lag between measurements^39,40^. Similarly, the backward in time cross-correlation function, 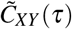, can be calculated. The key metric for non-equilibrium analysis is the average difference in cross-correlations between <u>the forward and backward</u> directions, denoted by 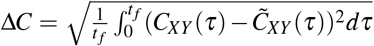. This quantity is calculated as the square root of the time integral over the forward and backward differences squared, integrated over a time window *t* _*f*_ . A non-zero value of Δ*C* signifies the presence of time irreversibility, implying that the system is not in detailed balance. Furthermore, the magnitude of Δ*C* reflects the strength of the flux arising from this time irreversibility. In essence, Δ*C* provides a quantitative measure of the degree of non-equilibrium within the system, which is directly linked to the extent of detailed balance breaking^34,39,40^.

We use the average difference Δ*C* for directly identifying dynamical phase transitions and bifurcations in cell fate decision-making systems by analyzing time series data. To maintain system stability during analysis and avoid unwanted transitions between steady states, we employ a relatively small diffusion coefficient. Time irreversibility, a key indicator of phase transitions, is then quantified by calculating the difference in cross-correlations between forward and backward time directions for each state: Δ*C*_*A*_ (system in state *S*_*A*_), Δ*C*_*O*_ (system in state *S*_*O*_), and Δ*C*_*B*_ (system in state*S*_*B*_). This strategy allows us to collect sufficient simulation data while keeping the system predominantly in the *S*_*A*_ state, minimizing the risk of transitions to *S*_*O*_ or *S*_*B*_. Notably, once the system switches to *S*_*O*_ or *S*_*B*_, Δ*C*_*A*_ loses its predictive power for the dynamical phase transition as the system has already exited state *S*_*A*_. The same argument holds true for evaluating Δ*C*_*B*_, Δ*C*_*O*_. As shown in Figure 4A with *b* = 0.21, all three metrics (Δ*C*_*A*_, Δ*C*_*O*_, and Δ*C*_*B*_) exhibit significant changes near the two bifurcation points, accompanied by small fluctuations.

**FIG. 4.**
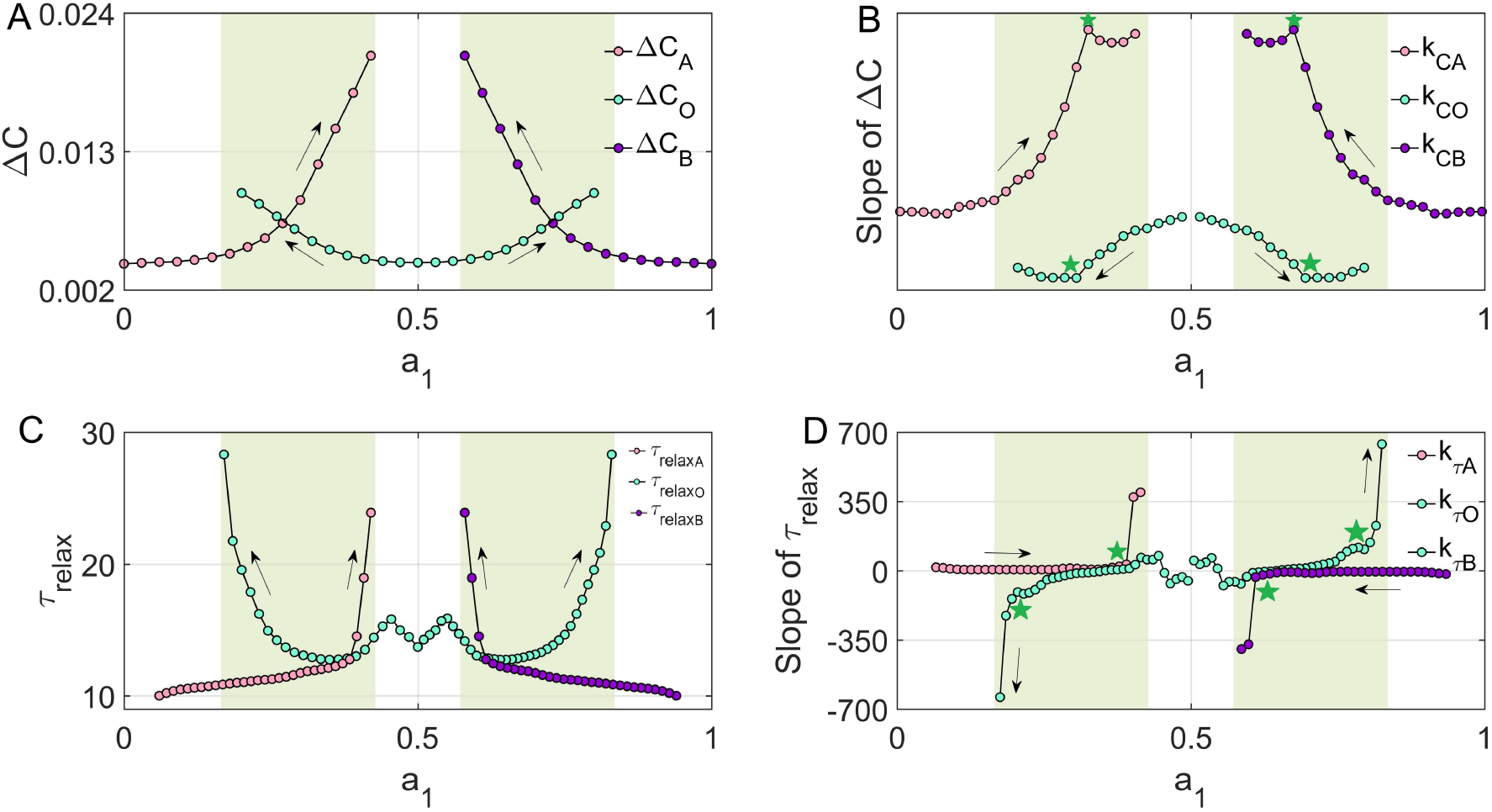
A: The average change of the forward and backward in time cross correlation function versus *a*_1_. B: The slope of Δ*C*) versus *a*_1_. C: The relaxation time of the autocorrelations *τ*_*relax*_ versus *a*_1_. D: The slope of the relaxation time *τ*_*relax*_ versus *a*_1_. The subscript *A* is for state *S*_*A*_, *B* is for state *S*_*B*_, *O* is for state *S*_*O*_. The parts with yellow-green background are the bistable region. (*b* = 0.21)

In Figure 4B, we present *k*_*CA*_ (the slope of Δ*C*_*A*_) for state *S*_*A*_, *k*_*CO*_ (the slope of Δ*C*_*O*_) for state *S*_*O*_, and *k*_*CB*_ (the slope of Δ*C*_*B*_) for state *S*_*B*_ plotted against *a*_1_. The data from simulations was fitted using the interpolation method to calculate the slopes of the lines. Arrows in the subfigure indicate the direction of parameter changes. Notably, *k*_*CA*_, *k*_*CO*_, and *k*_*CB*_ exhibit inflection points (green stars), indicative of significant changes nearing the four bifurcations. Particularly, the *S*_*O*_ state displays two inflection points, while both *S*_*A*_ and *S*_*B*_ states manifest an inflection point for each line, as *a*_1_ varies in both large and small directions (black arrows). Hence, the cross-correlation difference Δ*C* emerges as a pivotal early warning signal for potential transitions. Additionally, in 4C, we observe a sharp increase in relaxation time *τ*_*relaxA*_, *τ*_*relaxO*_, and *τ*_*relaxB*_ near the bifurcations, with minor fluctuations (*D* = 0.005). Furthermore, in 4D, we depict *k*_*τA*_, *k*_*τO*_, *k*_*τB*_ (the slope of the relaxation time *τ*_*relaxA*_, *τ*_*relaxO*_, *τ*_*relaxB*_ respectively) against *a*_1_. Remarkably, all slopes *k*_*τA*_, *k*_*τO*_, *k*_*τB*_ exhibit sharp increases (green stars), signaling significant increments in *τ*_*relax*_ as the bifurcations approach. These findings reveal that both the multi-dimensional information captured by Δ*C* and the simpler one-dimensional relaxation times present valuable tools for generating early warnings of phase transitions. This indicators are better for early warning detection, which are earlier than those of CSD (critical slowing down). This suggests that combining information from complex dynamical processes with simpler representations can provide robust methods for identifying critical transitions in cell fate decision-making systems.

### B. Direct Transdifferentiation

We explored the direct bifurcation of for cell transdifferentiation, specifically focusing on the mutual inhibition strength (*b* = 0.35) as depicted in Figure 5A. The phase diagram of two mutual inhibition and self-activation genes *X*_1_ and *X*_2_, as *a*_1_ increases, exhibits two-saddle-node bifurcations. Additionally, we present the three-dimensional potential landscapes with respect to *a*_1_, considering a diffusion coefficient of *D* = 0.005 and *b* = 0.35, as depicted in Figure 6A. Furthermore, we visualize the potential landscape *U* projected onto the line *X*_2_ = −*X*_1_ + 2 in Figure 6B, with the mutual inhibition strength set to *b* = 0.35. Additionally, Figure 6C illustrates a three-dimensional continuous potential landscape *U* versus *a*_1_ projected onto the line 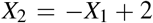, with a transformation represented by 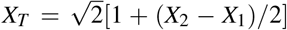. This representation also exhibits qualitative similarities to the marble-in-a-cup model. We observe two representative saddle-node bifurcations, indicative of first-order phase transitions. At lower self-activation strength (*a*_1_), a monostable differentiated cell state (*S*_*A*_) predominates. However, as *a*_1_ increases, a new stable differentiated cell state (*S*_*B*_) emerges through a saddle-node bifurcation. Subsequently, with further increments in *a*_1_, the stable differentiated cell state *S*_*A*_ undergoes a saddle-node bifurcation and vanishes. Notably, the transition of differentiated cells from state *S*_*A*_ to the new state *S*_*B*_ is facilitated by increasing *a*_1_ and decreasing self-activation strength *a*_2_. The valleys representing the differentiated state *S*_*A*_ disappear concomitant with the saddle-node bifurcation, elucidating the mechanism underlying cell type switching known as direct conversion transdifferentiation.

**FIG. 5.**
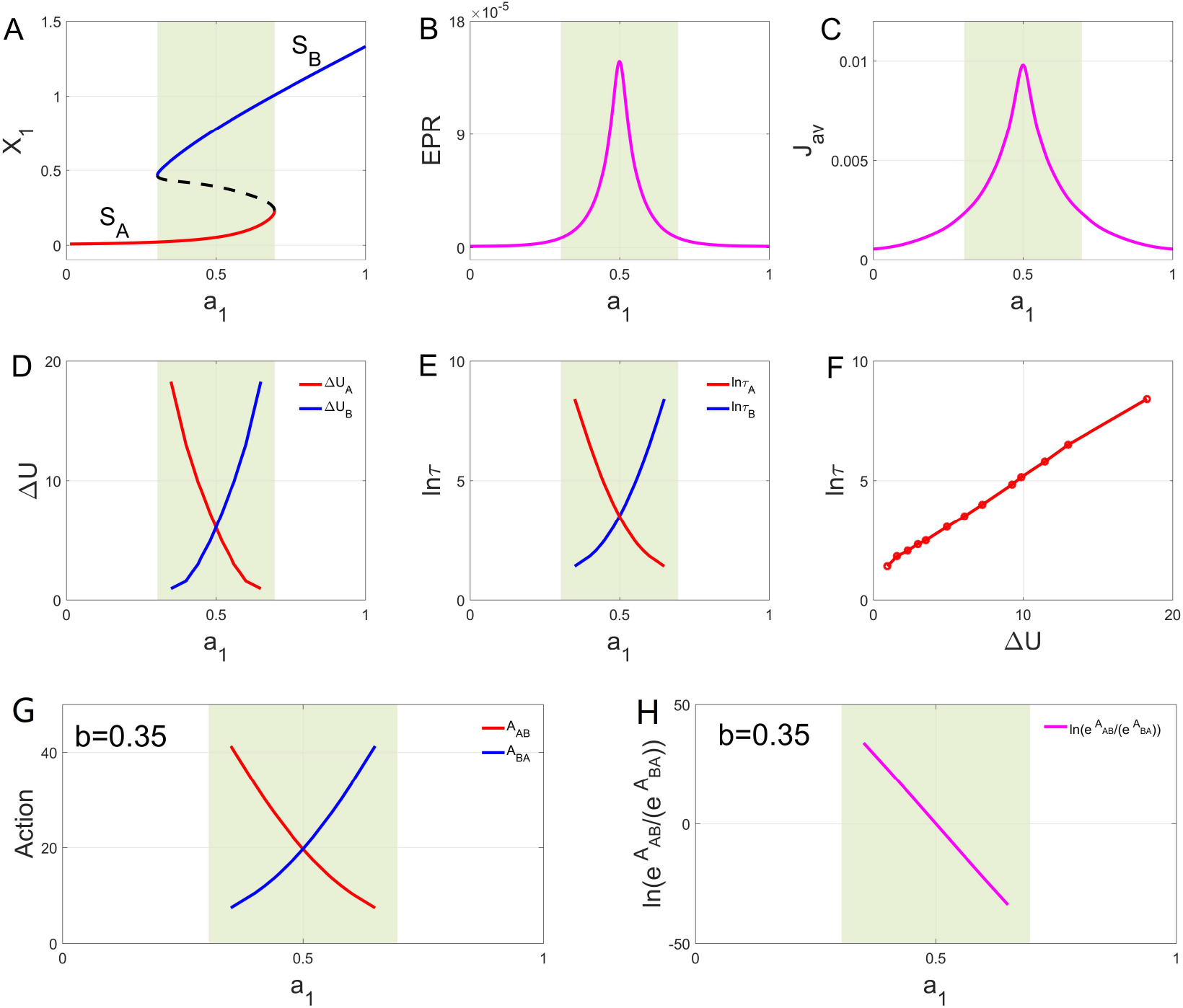
A: The phase diagram of two mutual inhibition and self-activation genes *X*_1_ and *X*_2_ with increasing the self-activation strength *a*_1_ giving arise to saddle node bifurcations (*b* = 0.35). B: The entropy production rate (*EPR*) versus *a*_1_. C: The average curl flux *J*_*av*_ versus *a*_1_. D: The barrier heights Δ*U* versus *a*_1_. E: The logarithm of the mean first passage time versus *a*_1_. F: The logarithm of the mean first passage time versus the barrier heights Δ*U* . G: The action *A*_*AB*_ of the probability of the dominant path from *S*_*A*_ to *S*_*O*_ and the action *A*_*BA*_ of the probability of the dominant path from *S*_*O*_ to *S*_*A*_ versus *a*_1_. H: The logarithm of the action *A*_*AB*_ divided by the action *A*_*BA*_ versus *a*_1_. The part with yellow-green background is the bistable region.

**FIG. 6.**
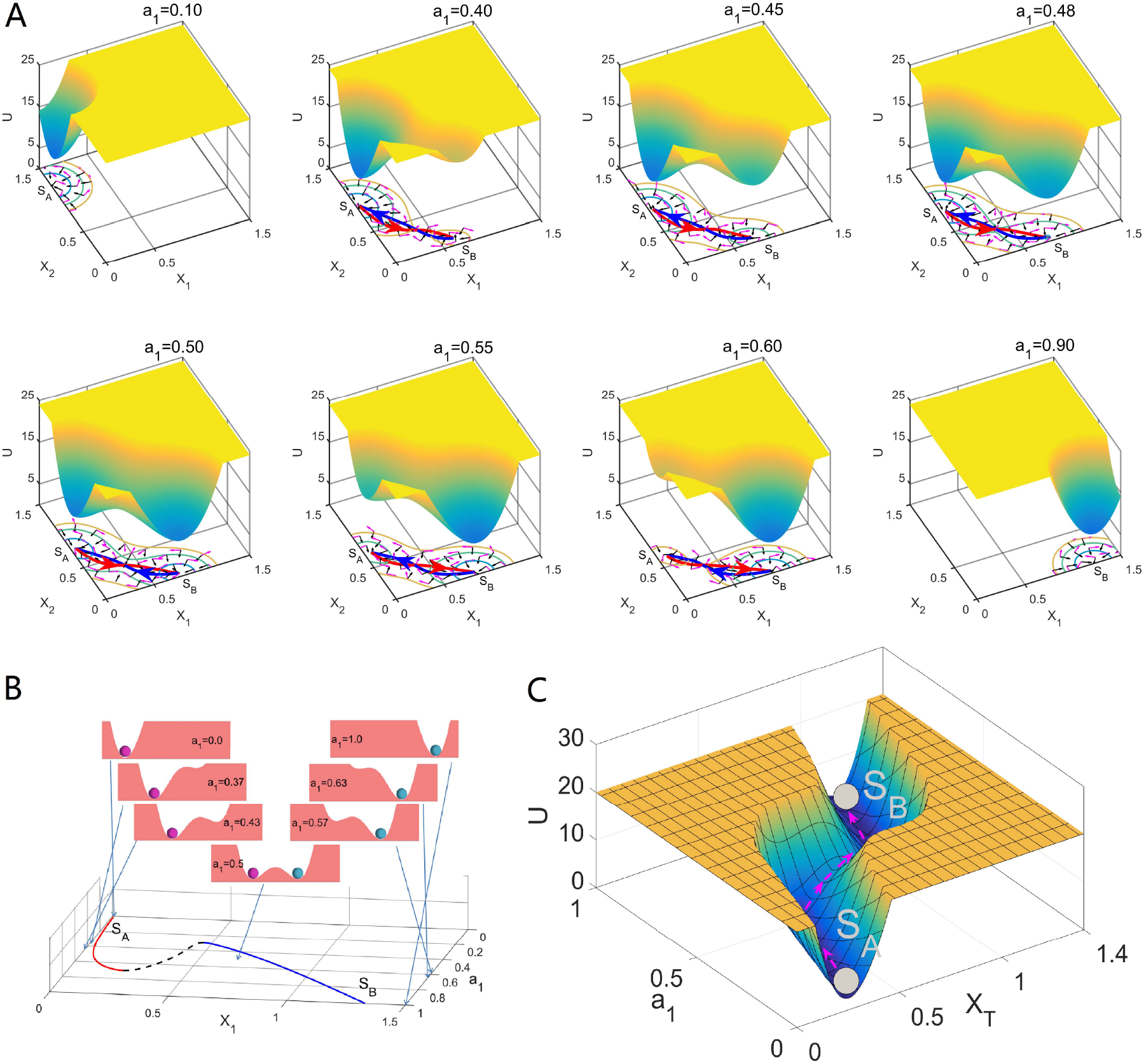
A: The potential landscapes versus *a*_1_ with the diffusion coefficient *D* = 0.005 (*b* = 0.35). The pathways on potential landscapes. The red lines represent the dominant population paths from the *S*_*A*_ state to the *S*_*B*_ state. The blue lines represent the dominant population paths from the *S*_*B*_ state to *S*_*A*_ state. B: The quantitative one-dimensional population potential landscapes by the projection versus *a*_1_ are on the top. The phase diagram is on the bottom. C: The continu<u>o</u>us potential landscape *U* versus the *a*_1_ and *X*_*T*_ with integral. The projection line is *X*_2_ = −*X*_1_ + 2 with the coordinate transformation 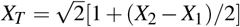.

Figure 5BC shows the entropy production rate (*EPR*) and the average curl flux *J*_*av*_ versus the self-activation strength *a*_1_ with the mutual inhibition strength *b* = 0.35. We can see that *EPR* and *J*_*av*_ have the same trends and both have one 0sharp peak, respectively. We show the barrier heights Δ*U* and the logarithm of the mean first passage time versus *a*_1_ in Figure 5DE. We show the logarithm of the mean first passage time versus the barrier heights Δ*U* in Figure 5F. Within the bistable region (highlighted by the yellow-green background), a critical point emerges where the barrier heights between states intersect. At this point, the weights of the two attractor states become equal (*a*_1_ = 0.5). This is the first order phase transition point which coincides with a wider distribution of the flux, leading to maximal values for both the average flux (*J*_*a*_*v*) and the entropy production rate (*EPR*). Furthermore, we also find the same correlation between the mean first passage time (kinetic switching time) and the barrier heights.

Figure 5G illustrates the actions (*A*_*AB*_ and *A*_*BA*_) linked to the dominant paths governing transitions between states *S*_*A*_ and *S*_*B*_. Here, *A*_*AB*_ denotes the action associated with the dominant path from *S*_*A*_ to *S*_*B*_, while *A*_*BA*_ represents the action for the dominant path from *S*_*B*_ to *S*_*A*_. Further exploration of this relationship is depicted in Figure 5H, showing the logarithm of the ratio between *A*_*AB*_ and *A*_*BA*_ as a function of the self-activation strength (*a*_1_). Our observations reveal an increasing trend in the logarithm of the dominant path probability from *S*_*A*_ to *S*_*B*_ relative to the dominant population path from *S*_*B*_ to *S*_*A*_ with increasing *a*_1_ (probability of the dominant path *P* ∼*e*^−*A*^). These findings collectively imply a shift in stability between states *S*_*A*_ and *S*_*B*_. As *a*_1_ increases, state *S*_*A*_ progressively loses stability, while state *S*_*B*_ gains stability. Consequently, transitions from *S*_*B*_ to *S*_*A*_ become increasingly challenging, while transitions from *S*_*A*_ to *S*_*B*_ become more accessible.

Our proposed non-equilibrium metrics, Δ*C*_*A*_ and Δ*C*_*B*_ (time irreversibility of cross-correlations for states *S*_*A*_ and *S*_*B*_), exhibit significant variations near the bifurcation points in Figure 7A. These variations are accompanied by minor fluctuations, suggesting heightened sensitivity around critical transitions. Similarly, the kinetic switching times, *k*_*CA*_ and *k*_*CB*_, depicted in Figure 7B, reveal inflection points (marked by green stars) near the bifurcations. Interestingly, both states *S*_*A*_ and *S*_*B*_ exhibit inflection points in their respective *k*_*C*_ curves as *a*_1_ is varied in both directions (indicated by black arrows).

**FIG. 7.**
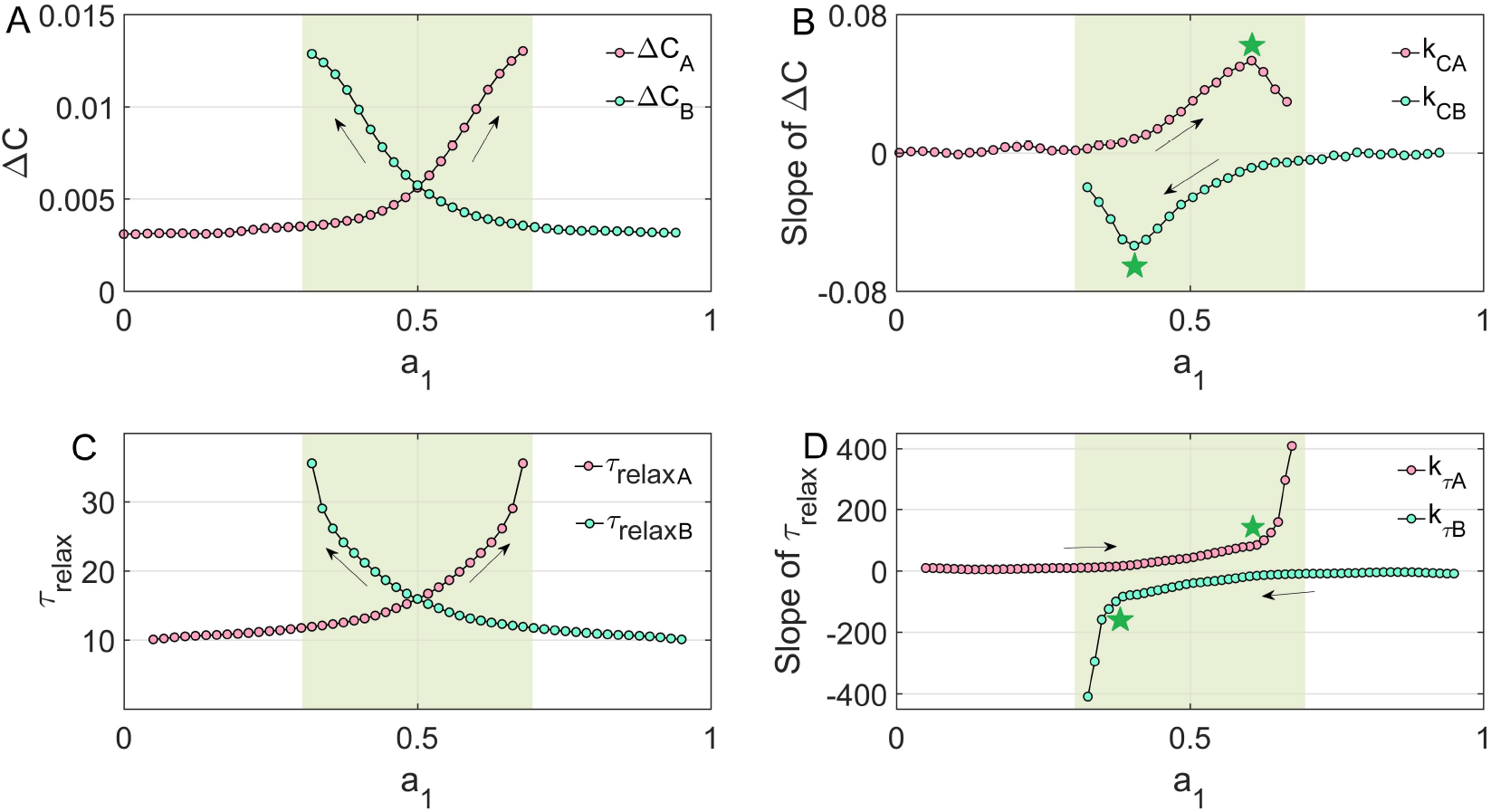
A: The average change of the forward and backward in time cross correlation function versus *a*_1_. B: The slope of Δ*C*) versus *a*_1_. C: The relaxation time of the autocorrelations *τ*_*relax*_ versus *a*_1_. D: The slope of the relaxation time *τ*_*relax*_ versus *a*_1_. The subscript *A* is for state *S*_*A*_, *B* is for state *S*_*B*_. The part with yellow-green background is the bistable region.(*b* = 0.35)

Furthermore, Figure 7C demonstrates a pronounced increase in the relaxation times, *τ*_*relaxA*_ and *τ*_*relaxB*_, for states *S*_*A*_ and *S*_*B*_, respectively, as the bifurcations are approached. These increases are accompanied by minor fluctuations (D = 0.005). Finally, the slopes of the relaxation times (*k*_*τA*_, *k*_*τB*_) plotted against *a*_1_ in Figure 7D show remarkable increases (marked by green stars). These sharp increases in slope signify significant enhancements in relaxation times as the system approaches critical transitions. We can find that the Δ*C*_*A*_ and Δ*C*_*B*_ are better early warning signals for predicting the dynamical phase transition than those obtained by CSD.

## IV. CONCLUSION

This study employs non-equilibrium landscape flux theory to explore into the intricacies of cell fate dynamics. Our investigation uncovers the pivotal role played by two key elements: the potential landscape and the curl flux. The landscape steers the system towards states of minimal potential energy, while the curl flux orchestrates transitions between these states, exerting influence on the likelihood of cell fate alterations.

Crucially, we demonstrate that the presence of non-zero curl flux initiates irreversible dominant pathways between multi-stable states, pathways not solely predictable through the potential landscape alone. This underscores the significance of non-equilibrium considerations in governing cell fate decision-making. Additionally, we pinpoint inflection points and significant slope changes in cross-correlation differences as precursory signals indicating impending bifurcations, offering opportunities for proactive intervention.

We further demonstrate the power of several quantitative measures for predicting cell fate transitions. These include the barrier height between states, the kinetic switching time (average first passage time), the entropy production rate, and the average flux. Interestingly, we observe a consistent trend between the average flux and entropy production rate, suggesting that both metrics offer a complementary perspective on the system’s stability and dynamics. Notably, the inherent rotational nature of the flux destabilizes attractor states, providing a dynamical origin for the observed phase transitions and dynamical bifurcations.

To predict critical transitions, we propose a set of non-equilibrium warning indicators: average flux, entropy production rate, and the time irreversibility of cross-correlations (Δ*C*). These indicators exhibit significant changes near bifurcations, enabling the early detection of transitions before the current state loses stability. Importantly, our approach provides earlier warnings compared to the traditional critical slowing down theory.

The ability to anticipate and manipulate cell fate decisions holds significant promise for stem cell applications. Our insights into non-equilibrium dynamics furnish valuable guidance for steering cell differentiation, reprogramming, and transdifferentiation. By discerning and regulating factors influencing curl flux and landscape topography, researchers can potentially navigate stem cell populations towards desired cellular destinies with heightened precision. For instance, minimizing non-equilibrium fluctuations or modulating potential landscapes through external stimuli could expedite controlled cell differentiation.

This study sheds light on the non-equilibrium forces governing cell fate decisions in active matter systems like stem cells. By unraveling the interplay among flux, potential landscape, and phase transitions, we establish a framework for not only understanding, but also potentially predicting and manipulating these processes. This opens doors for designing novel therapeutic strategies and advancements in regenerative medicine, allowing us to steer stem cell differentiation towards desired outcomes.

## ACKNOWLEDGMENTS

LX thanks supports by Natural Science Foundation of Jilin Province No. 20220101013JC and National Natural Science Foundation of China No.12234019.

